# Identification of a weight loss-associated causal eQTL in *MTIF3* and the effects of *MTIF3* deficiency on human adipocyte function

**DOI:** 10.1101/2022.10.16.512435

**Authors:** Mi Huang, Daniel Coral, Hamidreza Ardalani, Peter Spégel, Alham Saadat, Melina Claussnitzer, Hindrik Mulder, Paul W. Franks, Sebastian Kalamajski

**Affiliations:** Department of Clinical Sciences, Genetic and Molecular Epidemiology Unit, Clinical Research Centre, Lund University, Malmö, Sweden; Department of Nutrition, Harvard T.H. Chan School of Public Health, Boston, MA, USA; Department of Clinical Sciences, Unit of Molecular Metabolism, Clinical Research Centre, Lund University, Malmö, Sweden; Department of Chemistry, Centre for Analysis and Synthesis, Lund University, Lund, Sweden; Metabolism Program, Broad Institute of MIT and Harvard, Cambridge, USA

## Abstract

**Background:** Genetic variation at the *MTIF3* (Mitochondrial Translational Initiation Factor 3) locus has been robustly associated with obesity in humans, but the functional basis behind this association is not known.

**Methods:** Here, we applied luciferase reporter assay to map potential functional variants in the haplotype block tagged by rs1885988 and used CRISPR-Cas9 to edit the potential functional variants to confirm the regulatory effects on *MTIF3* expression. We further conducted functional studies on MTIF3-deficient differentiated human white adipocyte cell line (hWAs-iCas9), generated through inducible expression of CRISPR-Cas9 combined with delivery of synthetic *MTIF3*-targeting guide RNA.

**Results:** We demonstrate that rs67785913-centered DNA fragment (in LD with rs1885988, r^2^>0.8) enhances transcription in a luciferase reporter assay, and CRISPR/Cas9 edited rs67785913 CTCT cells show significantly higher *MTIF3* expression than rs67785913 CT cells. Perturbed *MTIF3* expression changed the expression of mitochondrial DNA-encoded genes, and reduced mitochondrial respiration, as well as altered endogenous fatty acid oxidation. Furthermore, after glucose restriction, the *MTIF3* knockout cells retained more triglycerides than control cells.

**Conclusions:** This study demonstrates an adipocyte function-specific role of *MTIF3*, which originates in the maintenance of mitochondrial function, providing potential explanations for why *MTIF3* genetic variation at rs67785913 is associated with body corpulence and response to weight loss interventions.

## Introduction

Over 650 million people are obese and often suffer from metabolic abnormalities, including dyslipidemia, type 2 diabetes, and hypertension (1, 2). It is widely believed that obesity results from an interplay between genetic and environmental factors (3), but the biological mechanisms behind these interactions are poorly understood.

Genetic variation (rs12016871) at *MTIF3* (encoding the Mitochondrial Translation Initiation Factor 3 protein (4)) has been robustly associated with body mass index (BMI) in humans (5). Several subsequent studies have linked *MTIF3* genetic variation with the response to weight loss interventions, including diet, exercise and bariatric (6, 7) and with weight-related effects of habitual diet (8). For example, analyses in two of the world’s largest randomized controlled weight loss trials (Diabetes Prevention Program [DPP] and Look AHEAD) found that homozygous minor allele carriers (rs1885988) were slightly more prone to weight gain in the control arm, yet achieved significantly greater weight loss at 12-month post-randomization and retained lost weight longer (18-36 months) than major allele carriers (6). Elsewhere, the same locus has been associated with greater and more sustained weight loss following bariatric surgery (7).

Mtif3 loss in the mouse results in cardiomyopathy owing to impaired translation initiation from mitochondrial mRNAs and uncoordinated assembly of OXPHOS complexes in heart and skeletal muscle (9). In the human hepatocyte-like HepG2 cell line, MTIF3 loss decreases the translation of the mitochondrial-encoded ATP synthase membrane subunit 6 (*ATP6*) mRNA without affecting cellular proliferation (10). No human genomic mutations leading to total MTIF3 deficiency have been reported, but the studies outlined above suggest that *MTIF3* may influence obesity predisposition and weight loss potential by modulating mitochondrial function; thus, *MTIF3* may play a key role in adipose tissue metabolic homeostasis, as adipocyte mitochondria not only provide ATP, but also impact adipocyte-specific biological processes such as adipogenesis, lipid metabolism, thermogenesis, and regulation of whole-body energy homeostasis (11, 12).

The purpose of this study was to fine map the causal DNA variations behind genetic variations in *MTIF3* and diet interactions on weight change. Second, we assessed whether MTIF3 loss in human white adipocytes influences adipocyte-specific, obesity-related traits under basal and perturbed metabolic conditions. For the latter, we used glucose restriction to mimic the effects of *in vivo* lifestyle interventions focused on energy restriction and energy expenditure.

## Results

### rs67785913 is a regulatory variant for *MTIF3* expression

The *MTIF3* rs1885988 C allele is associated with enhanced weight loss and weight retention following lifestyle intervention trials in DPP and Look AHEAD cohorts (13). In the GTEx database, the rs1885988 associates with an eQTL in subcutaneous fat (Figure 1A), with C allele carriers having significantly higher *MTIF3* expression (normalized effect size: 0.15, p = 0.0000032).

**Figure 1.**
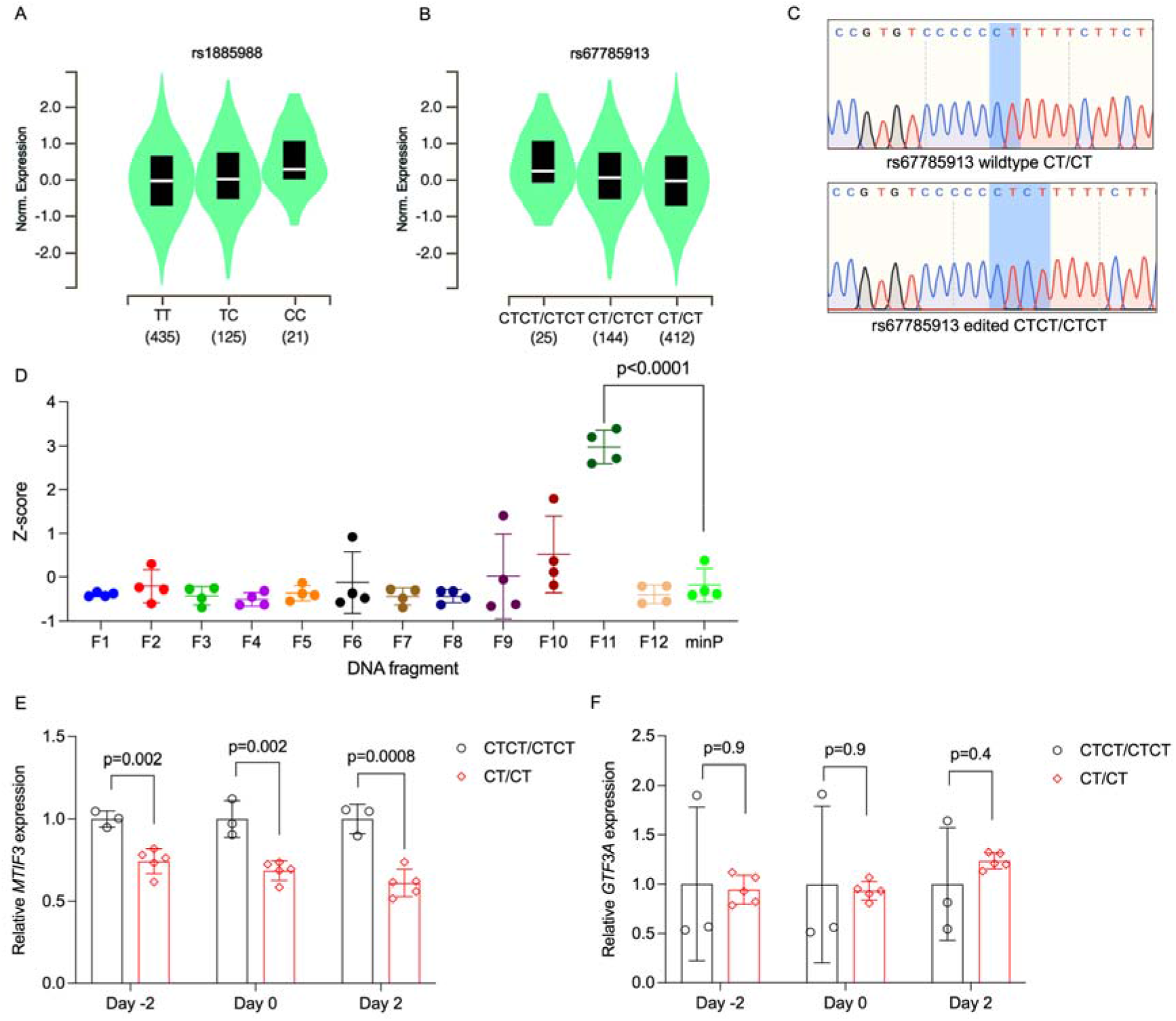
**A.** Violin plot of *MTIF3* expression in subcutaneous adipose tissue for rs1885988 from Genotype-Tissue Expression (GTEx) Project eQTL. **B.** Same as in **A**, but for rs67785913. **C.** Normalized Z-score plot of luciferase reporter assays using vectors carrying different DNA fragments of the *MTIF3* gene cloned into pGL4.23 luciferase reporter vector. Hypothesis testing was performed by comparing the transcriptional enhancer activity of each of the 12 vectors (F1-12) to the empty vector (minP). All data were plotted as mean± SD, n = 4 independent experiments, p values are presented in each graph; ordinary one-way ANOVA was used for statistical analysis. **D.** Representative Sanger sequencing traces of rs67785913 CTCT/CTCT and CT/CT clones obtained after CRISPR/Cas9-mediated allele editing and single cell cloning. **E.** Relative *MTIF3* expression in rs67785913 allele-edited cells two days before, at, or two days post-differentiation induction (day −2, 0, 2, respectively). n = 3 clonal populations for CTCT/CTCT genotype, n = 5 clonal populations for CT/CT genotype, error bars show standard deviation. **F.** as in **E**, but for *GTF3A* expression. Two-tailed Student’s t test was used; p values are presented in each graph.

To experimentally validate and fine map the potential causal DNA variation in the haplotype block tagged by rs1885988, we looked up all tightly linked (r^2^>0.8) SNPs in HaploReg database v4.1(14). We then PCR-amplified and cloned 12 DNA fragments from that haploblock, altogether comprising the linked SNP loci, into luciferase reporter plasmids. As shown in Figure 1D, by comparing the luciferase signals with minimal promoter (minP) construct, only one DNA fragment (F11), encompassing the rs67785913 locus, could enhance luciferase transcription. Coincidentally, the rs67785913 also shows an eQTL effect on *MTIF3* expression in subcutaneous adipose tissue in GTEx database, with the major CT allele associated with lower expression than the minor CTCT allele (normalized effect size: −0.16, p = 3.0 × 10^−8^) (Figure 1B). To demonstrate an allele-specific regulatory effect on *MTIF3* expression, we then used CRISPR-Cas9 to substitute the major CT for the minor CTCT allele at the rs67785913 locus in the preadipocyte hWAs cell line. Due to rather low CRISPR editing efficiency of that locus, we needed to genotype over 700 single cell clones to obtain five CT/CT and three CTCT/CTCT clones without random indels, as confirmed by Sanger sequencing (Figure 1C). We then examined *MTIF3* expression in these clones at pre- and post-adipogenic differentiation induction, and found rs67785913 CTCT/CTCT to confer higher *MTIF3* expression at all time points (Figure 1E). As rs67785913 also correlates with an altered *GTF3A* expression in other tissues (e.g. muscle, lung), we also detected, but found no apparent difference in *GTF3A* expression in rs67785913 edited cells (Figure 1F).

### Generating inducible Cas9-expressing pre-adipocyte cell line (hWas-iCas9)

In pre-adipocytes, marginally different passage numbers between control and experimental groups during adipogenic differentiation can cause confounding. This can originate during work to create genetic knockouts/knockins and precludes meaningful studies of gene x environment interactions, where it is often desirable to use similarly differentiated cells with comparable baselines for e.g. triglyceride or mitochondrial content. Unfortunately, in the above-described rs67785913 allele-edited cells, this problem appeared and precluded further proper functional genomics work. To circumvent this, we established an inducible Cas9-expressing pre-adipocyte cell line, that allowed us to model an eQTL by inducing genetic knockdown in differentiated hWAs cells. As illustrated in Figure 2, we co-transfected hWAs with two plasmids: one encoding piggyBac transposase, and the other carrying piggyBac transposon-flanked doxycycline-inducible Cas9 and constitutively expressed puromycin resistance genes. In this setup, piggyBac transposase drives the integration of the piggyBac-transposon flanked genes, and transgenic cells are then selected and expanded in puromycin-supplemented culture medium. We have thus obtained an hWAs cell line with doxycycline-inducible Cas9 expression and maintained adipogenic differentiation capacity (henceforth called hWAs-iCas9). The Cas9-expressing differentiated cells could then be transfected with relatively low molecular weight synthetic single guide RNAs (sgRNAs) that complex with intracellularly expressed Cas9 and target the gene exon of interest to generate random indels (in essence, gene knockouts). We used this method here to determine the functional role of *MTIF3* in adipocyte biology.

**Figure 2.**
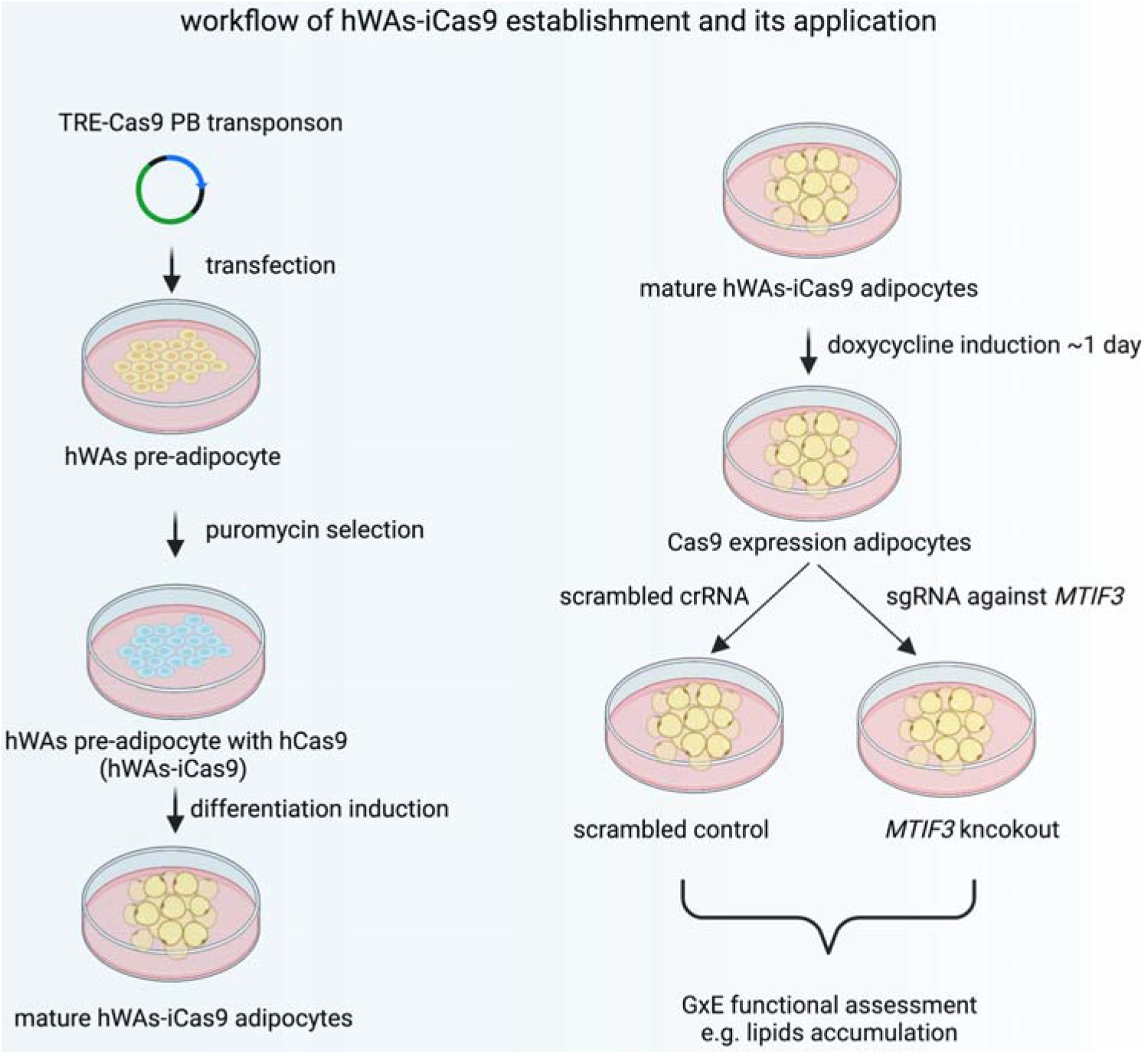
The workflow of establishing hWAs-iCas9 cell line and its application in studying MTIF3 and environment interactions in *vitro*.

### Generation of *MTIF3* knockout in hWAs-iCas9 adipocytes

To investigate the role of *MTIF3* in human adipocyte development and energy metabolism we generated stable *MTIF3* knockouts in differentiated hWAs-iCas9 adipocytes. We designed Cas9-specific sgRNA to generate random indels in the exon expressed in all three *MTIF3* protein-encoding transcripts (Figure 3A) and obtained a >80% reduction in MTIF3 protein levels in every experiment, as assessed by Western blotting (Figure 3D and E). To assess off-target effects of CRISPR-Cas9, we also performed T7EI assays on PCR-amplified top 5 predicted off-target sites and did not observe any detectable off-targeting (data is not shown).

**Figure 3.**
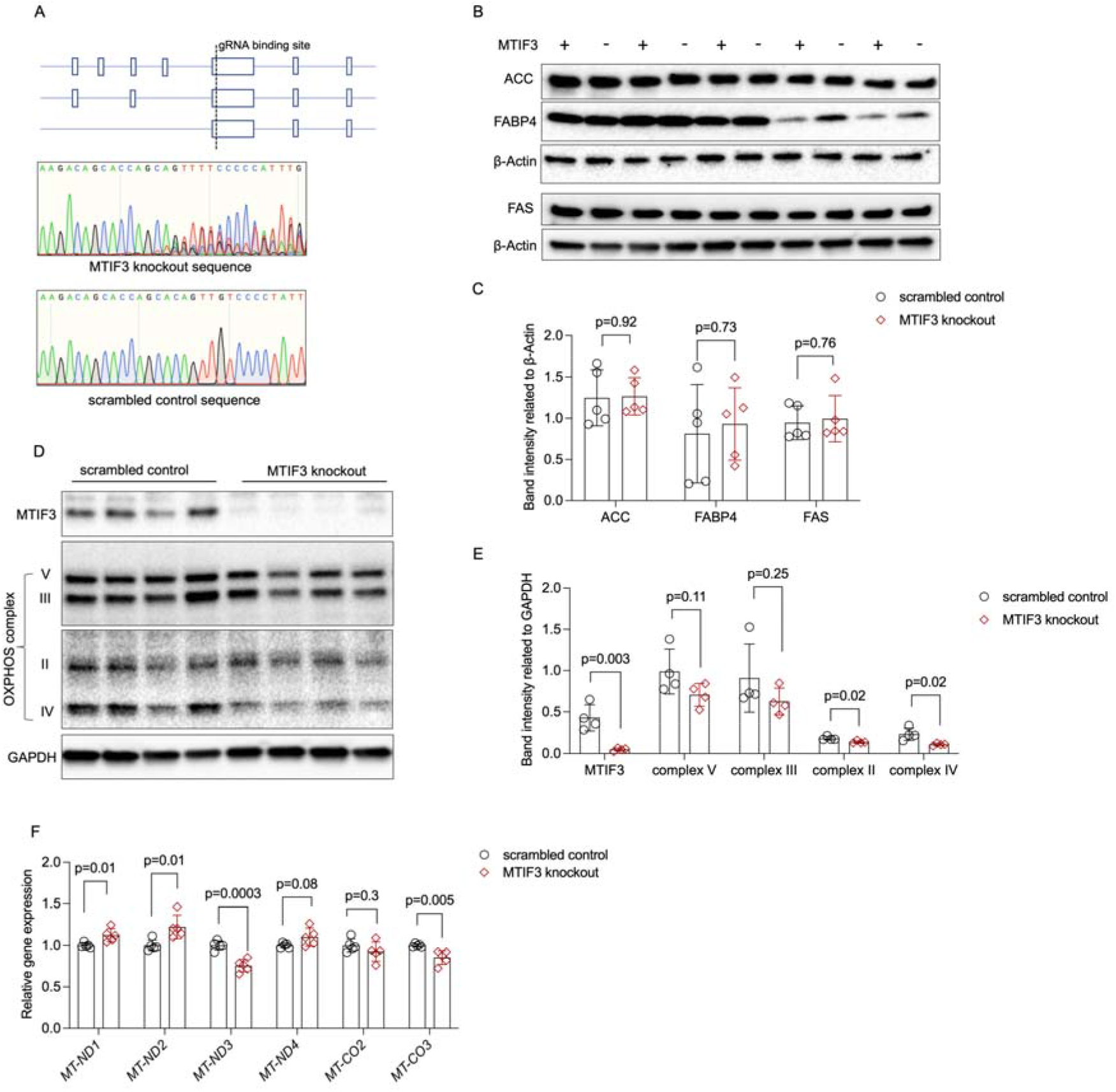
MTIF3 perturbation in mature adipocytes does not affect adipocyte-specific protein expression but disrupts mitochondrial OXPHOS complex expression. **A.** An illustration of Cas9-specific sgRNA binding site in the exon expressed in all three *MTIF3* protein-encoding transcripts and representative Sanger sequencing of control and knockout hWAs mature adipocytes. **B.** Immunoblots of adipocyte markers in scrambled control and *MTIF3* knockout adipocytes, n = 5 independent experiments. **C.** Quantitative analysis of the band densities in **B. D.** Immunoblots of MTIF3 and mitochondrial OXPHOS complexes in scrambled control and *MTIF3* knockout mature adipocytes, n = 4 independent experiments. **E.** Quantitative analysis of the band densities of the proteins in **D**. **F.** qPCR for mitochondrial gene expression in scrambled control and MTIF3 knockout adipocytes, n = 5 independent experiments. Error bars show standard deviation in all plots. Statistical analysis was performed using two-tailed Student’s t test, p values are presented in each graph.

### *MTIF3* knockout decreases mitochondrial OXPHOS complex expression IV content, and disrupts mitochondrial DNA-encoding gene expression, without affecting adipogenic differentiation marker expression in hWAs-iCas9 adipocytes

MTIF3 is a mitochondrial translation initiation factor; thus, we examined the effects of MTIF3 ablation on differentiated hWAs adipocyte mitochondrial respiration chain. Assessed by western blotting on *MTIF3* knockout vs. control cell lysates, OXPHOS complex IV decreased by 54% (n = 4, p = 0.02), while complex III and V nominally decreased (Figure 3D and E). OXPHOS complex II appears to be decreased by 24% in *MTIF3* knockout cells (n = 4, p = 0.02). In contrast, the white adipocyte-specific markers ACC, FABP4 and FAS were unaffected by the MTIF3 perturbation in white adipocytes (Figure 3B and C). Morever, using qPCR, we observed an altered expression of several mitochondrial DNA-encoding genes in *MTIF3* knockout cells. Specifically, MTIF3 deficiency led to higher expression of *MT-ND1* (n = 5, p = 0.01), *MT-ND2* (n = 5, p = 0.01), a trending increase of *MT-ND4* (n = 5, p = 0.08), and lower expression of *MT-ND3* (n = 5, p = 0.0003), and *MT-CO3* (n = 5, p = 0.005).

### *MTIF3* knockout affects mitochondrial respiration in hWAs-iCas9 adipocytes

To investigate mitochondrial function in MTIF3-ablated differentiated hWAs adipocytes, we used the Seahorse Mito Stress Test to measure the OCR. Additionally, to avoid potential cofounders caused by the high glucose content in the differentiation medium, we adapted the cells into 1g/L growth medium for three days before running this assay. As shown in Figure 3, *MTIF3* knockout cells exhibited lower basal OCR, as well as lower ATP-forming capacity, the latter estimated by calculating OCR decrease after blocking ATP synthase with oligomycin (Figure 4A, B and C). MTIF3 knockout cells also showed a trending decrease on maximal respiration OCR (Figure 4D, p = 0.07). Furthermore, both *MTIF3* knockout and control cells, had comparable proton leak OCR, non-mitochondrial respiration OCR and coupling efficiency (Figure 4E, F and G).

**Figure 4.**
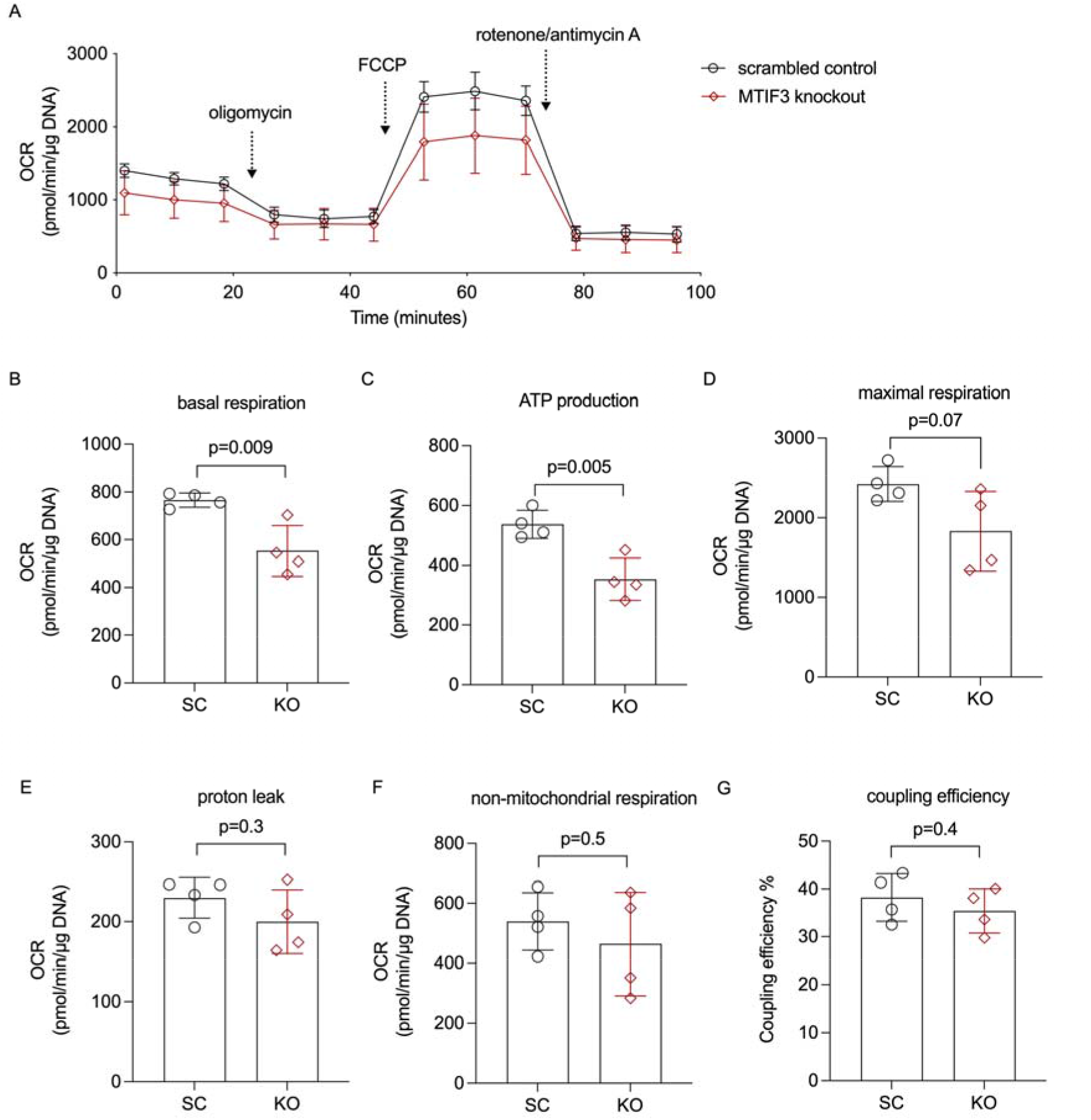
Cellular mitochondrial respiration in hWAs adipocytes. **A.** The average oxygen consumption rate (OCR) traces during basal respiration, and after addition of oligomycin, FCCP, and rotenone/antimycin A. **B.** Basal respiration OCR, n = 4 different cell passages. **C.** ATP production OCR, n = 4 different cell passages. **D.** Maximal respiration OCR, n = 4 different cell passages. **E**. Non-mitochondrial respiration OCR, n = 4 different cell passages. **F.** Proton leak OCR, n = 4 different cell passages. **G.** Coupling efficiency, n = 4 different cell passages. Error bars show standard deviation. Statistical analyses were performed using paired Student’s t test in each condition, p values are presented in each graph.

### *MTIF3* knockout affects hWAs-iCas9 adipocyte endogenous fatty acid oxidation

Next, we performed Seahorse assays to compare the endogenous fatty acid oxidation in *MTIF3* knockout vs. control cells, treated with etomoxir (an inhibitor of carnitine palmitoyl transferase). We found that MTIF3 ablation mimics the effect of etomoxir on basal endogenous fatty acid oxidation OCR. Furthermore, while etomoxir decreases basal fatty acid oxidation OCR in control cells, it does not markedly decrease it in *MTIF3* knockout cells (Figure 5A and B).

**Figure 5.**
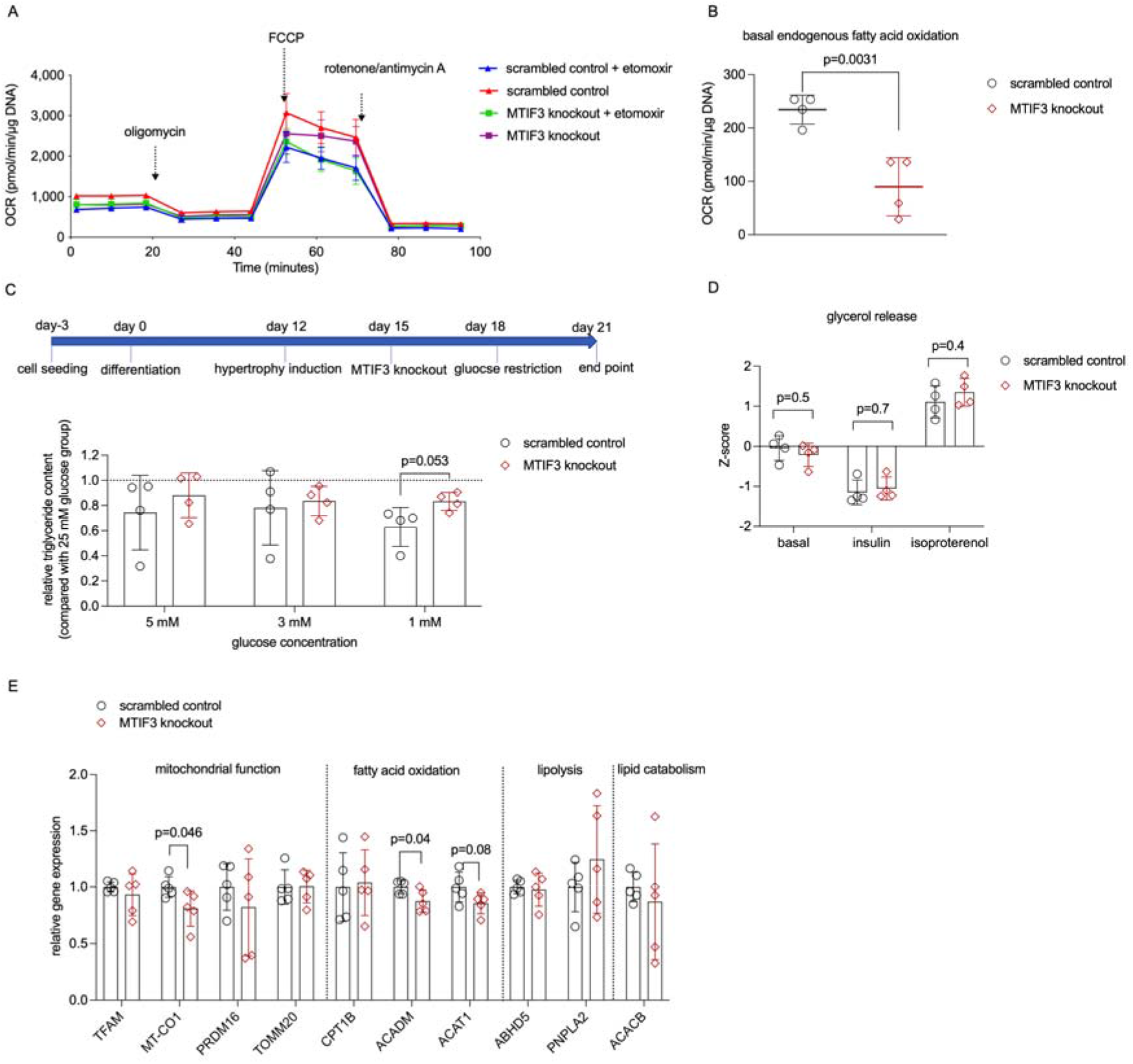
MTIF3 perturbation affects adipocyte fatty acid oxidation. **A.** A representative Seahorse OCR trace for endogenous fatty acid oxidation assay. *MTIF3* knockout and scrambled control adipocytes were treated with or without etomoxir for 15 minutes before the assay. Following the basal OCR measurement, oligomycin, FCCP (carbonyl cyanide-p-trifluoromethoxyphenylhydrazone), and rotenone + antimycin A were added sequentially to measure the detection of ATP production OCR, maximal respiration OCR and non-mitochondrial respiration OCR. **B.** Basal endogenous fatty acid oxidation oxygen consumption rate in scrambled control and *MTIF3* knockout adipocytes, n = 4 independent experiments. **C.** Total triglyceride contents in scrambled control and *MTIF3* knockout adipocytes at normal feeding condition and glucose restricted conditions, n = 4 independent experiments. **D.** Z-score-normalized data for glycerol release in scrambled control and MTIF3 knockout adipocytes under basal, insulin-stimulated, and isoproterenol-stimulated conditions, n = 4 independent experiments. **E.** qPCR for mitochondrial and adipocyte-related gene expression in scrambled control and *MTIF3* knockout adipocytes. Error bars show standard deviation in all plots. Statistical analysis was performed using two-tailed Student’s t test, p values are presented in each graph.

### *MTIF3* knockout affects hWAs-iCas9 adipocyte triglyceride content after glucose restriction challenge

To mimic the interactions between MTIF3 content and dietary intervention on weight change, we generated hypertrophic control and *MTIF3* knockout hWAs-iCas9 adipocytes and then used glucose-limited medium to mimic energy restriction *in vivo* (schematic shown in Figure 5C). Triglyceride content decreased both in control and *MTIF3* knockout cells after 3 days of different levels of glucose restriction when compared with normal feeding (25 mM glucose) medium. Interestingly, a more extensive decrease in triglyceride content occurred in control cells cultured in 1 mM glucose medium (p = 0.053, n = 4), and a similar trending decrease, albeit with higher coefficient of variation, occurred in 3 and 5 mM glucose medium (Figure 5C).

### *MTIF3* knockout does not affect hWAs adipocyte lipolysis

To examine the effects of *MTIF3* knockout on lipolysis, we measured basal, insulin-attenuated, and isoproterenol-stimulated glycerol release in differentiated hWAs cells. As shown in Figure 5D, in all three conditions, glycerol release in control and *MTIF3* knockout cells was comparable.

### *MTIF3* knockout affects mitochondrial function- and fatty acid oxidation-related gene expression

Next, we examined how MTIF3 ablation affects the gene expression programmes pertinent to mitochondrial function, fatty acid oxidation, lipolysis and lipid catabolism. As shown in Figure 5E, *MTIF3* knockout cells had decreased expression of the mitochondria-related *MT-CO1*, and the fatty acid oxidation-related *ACADM* and *ACAT1*, but unchanged expression of other genes involved in mitochondrial function and lipid metabolism (*TFAM, TOMM20, PRDM16, CPT1B, ABHD5, PNPLA2, ACACB*).

### *MTIF3* knockout results in glucose-level-depending alterations in metabolism

The combined GC/MS and LC/MS metabolite profiling resulted in relative quantification of 110 metabolites. First, we analyzed metabolite profiles at a global level using PCA. The score plot reveals a clear systematic difference in the metabolite profile between cells on normal feeding and glucose restriction (Figure 6A). Interestingly, differences between MTIF3 and control cells at normal feeding are observed along principal component 1 (PC1), whereas differences between genotypes at glucose restriction are observed along PC2, suggesting the effect of MTIF3 ablation to depend on the calorie level. Next, to identify alterations in metabolite levels underlying this differential response, we analysed data using orthogonal projections to latent structures discriminant analysis (OPLS-DA) separately at normal feeding (two components R2 = 0.82, Q2 = 0.66) and at glucose restriction (two components, R2 = 0.95, Q2 = 0.52). These analyses revealed systematic differences between genotypes at both growth conditions (Figure 6B and C). Next, to examine whether the differences between genotype depended on growth condition, we combined the correlations from the two OPLS-DA models into a shared and unique structures plot (Figure 6D). These analyses revealed levels of intermediates in cytosolic metabolic pathways connected to the glycolysis, such as glycerate 3-phosphate, glycerol 2-phosphate, UDP-N-acetylglucosamine and ribose 5-phosphate, to be lower in *MTIF3* knockout cells at both feeding conditions. Interestingly, levels of fatty acids, ranging from nine to 17 carbons and including several odd-chain fatty acids, were lower in the *MTIF3* knockout cells only at normal feeding. At glucose-restricted conditions, levels of both essential and non-essential amino acids were lower in the knockout. Finally, we analysed data using two-way ANOVA, incorporating feeding condition and genotype, thereby providing information on effects at the individual metabolite level. These analyses revealed 18 and 20 significantly different metabolites between control and *MTIF3* knockout cells at normal feeding and glucose restricted conditions, respectively (*q* < 0.05). These included ribose 5-phosphate, glycerate 3-phosphate, glycerol 2-phosphate and, glycerol 3-phosphate (Figure 6E).

**Figure 6.**
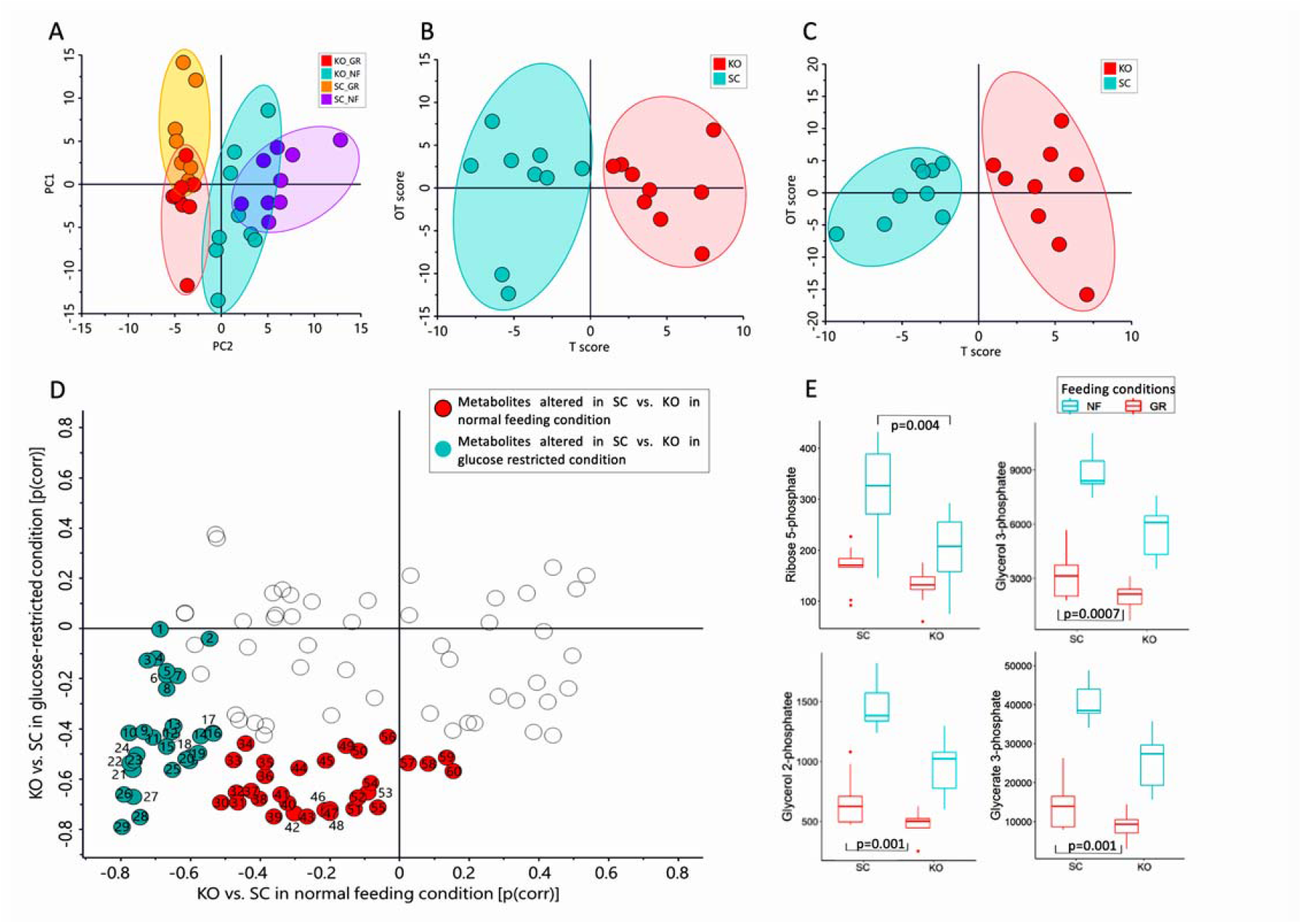
Mass spectrometry-based metabolomics data for control (SC) and *MTIF3* knockout (KO) cells in normal (NF) and glucose-restricted (GR) conditions. **A.** Principal component analysis (PCA) score plot displaying the discrimination between *MTIF3* knockout and control cells in normal and glucose-restricted conditions (PC1: 28%, PC2: 19%). **B.** Orthogonal projections to latent structures discriminant analysis (OPLS-DA) score plot showing classification of *MTIF3* knockout and control cells in normal feeding condition. **C.** OPLS-DA score plots showing classification of *MTIF3* knockout and control cells in glucose-restricted condition. **D.** Shared and unique structures (SUS) plot, based on OPLS-DA models in (**B**) and (**C**), showing feeding condition-dependent differences between MTIF3 knockout and control cells. **E.** Box plots showing the abundance of some of the significantly altered metabolites in normal and *MTIF3* knockout cells in normal and glucose-restricted conditions. Statistical analysis was performed using Two-way ANOVA test, p values are presented in each graph.

## Discussion

Excessive weight gain caused by dietary excess, and its effects on adipocyte lipid metabolism, can cause life-threatening disease (16–19). Findings from clinical trials (13), a bariatric surgery case series (7) and epidemiological cohorts (8) showed the *MTIF3* variation modulates weight loss-promoting exposures on body weight (see also UK Biobank analysis in Supplemental Table 2). Here, we validated *MTIF3* rs1885988 C allele correlates with higher *MTIF3* expression in subcutaneous fat tissue, and our *in vitro* luciferase reporter assay and CRISPR-Cas9 genome editing results revealed that the tightly linked rs67785913 variant is likely to be the actual eQTL for *MTIF3* expression (Figure 1).

Pre-adipocyte cell lines have been used extensively for studying adipogenic differentiation (20), but their application for lipid metabolism studies has been limited, especially in the context of gene-environment interaction. This is largely due to the variation of differentiation capacity across cell passages (21) and differential genetic effects on adipocyte differentiation (22), which can alter baseline phenotypes in differentiated cells. Therefore, we established an inducible Cas9-expressing human pre-adipocyte cell line (hWAs-iCas9), which enabled us to generate gene knockout of interest in pre-differentiated adipocytes, thus circumventing these limitations (Figure 2). Using the inducible knockout cell model, we then tested interactions between *MTIF3* and environmental changes (“lifestyle mimetics”). Our experiments in white adipocytes revealed *MTIF3* knockout-mediated disruption of mitochondrial respiration owing to decreased OXPHOS complex expression. This led to perturbed cellular functions, including reduced fatty acid oxidation and a trending increase in intracellular triglyceride content. These data indicate that *MTIF3* plays an important role in lipid metabolism in human adipocytes.

In adipose tissue, mitochondria play an essential role not only by ensuring ATP supply but also by triggering cellular signalling pathways that require reactive oxygen species (ROS) generated by OXPHOS complex I and III (23). These, and complex II, IV, and V, are partially encoded by mitochondrial DNA (24). The post-transcriptional rate-limiting translation step can be promoted by MTIF3, as it facilitates initiation complex formation on mitochondrial 55S ribosomes in the presence of MTIF2, fMet-tRNA and poly(A,U,G) (25). Previous studies have shown that loss of MTIF3 results in an imbalanced assembly of OXPHOS complexes in muscle and heart in mouse (9) and decreased translation of ATP6 mRNA in hepatocyte-like HepG2 cells (10). Our data showed that MTIF3 deficiency in adipocytes results in lower content of mitochondrial complex IV and altered several mitochondrial DNA-encoding genes, along with impaired mitochondrial respiration rate (Figure 4). Altogether, our results suggest that MTIF3 vastly affects the mitochondrial electron transport chain.

Mitochondrial dysfunction in white adipose tissue has been frequently associated with obesity (26), with the presumed mechanisms being decreased fatty acid oxidation and ATP production (reviewed in (27)). We have found that one causal link connecting these factors may be the MTIF3 content (Figure 4 and 5). Furthermore, a previous study described an inverse relationship between mitochondrial capacity and weight change (28); thus, conceivably, any genetic variation-modulated change in *MTIF3* expression could influence a dietary intervention outcome. We attempted to test this hypothesis *in vitro*, exposing hypertrophic adipocytes to glucose restriction, thereby mimicking weight loss-promoting exposures *in vivo*. We found that MTIF3-deficient adipocytes exposed to glucose restriction challenge responded less through changed triglyceride content, indicating limited capacity for lipid metabolism under glucose restriction (Figure 5). One should bear in mind that our data were generated *in vitro* in cells highly depleted of MTIF3. At this point, we do not know to which extent a more moderate MTIF3 deficiency *in vivo*, e.g. conferred by common genetic variants, will influence adipogenic differentiation and long-term diet-induced weight loss in a controlled study.

Finally, the metabolomic analysis added another dimension to the role of MTIF3 in regulating adipocyte metabolism (Figure 6). The lower content of glycolysis intermediates and odd chain fatty acids in *MTIF3* knockout adipocytes could indicate blunted lipogenesis, while the decreased essential and non-essential amino acid level in *MTIF3* knockout cells under glucose restriction could be an adaption to the lower energy supply owing to impaired mitochondrial function (40). The metabolite data, and the earlier described lower fatty acid oxidation capacity of MTIF3-deficient cells, suggest that MTIF3 plays a vital role in triglyceride metabolism in adipocytes, and provides further insight into the previously reported role of this protein on weight loss induced by dietary intervention (13).

In summary, we observed rs67785913 is the functional variant that causally modulates *MTIF3* expression. We also established a novel, efficient, and simple method to make *MTIF3* knockout in differentiated adipocyte cell line, and it should be generalizable for making other gene knockouts. We found that MTIF3 is essential for proper synthesis of protein complexes in adipocyte mitochondrial electron transport chain, with consequential effects on mitochondrial function and lipid metabolism. MTIF3 content can also affect the outcome of interactions with exposures related to energy restriction and expenditure. Altogether, this helps to explain the association between *MTIF3* genetic variation and several human body weight-related variables and may suggest that people carrying cis-eQTLs that increase *MTIF3* expression could benefit more from lifestyle intervention aiming at body weight control.

## Research Design and Methods

### Allele-specific *MTIF3* expression in subcutaneous adipose tissue

Data and plots presented in this manuscript for allele-specific *MTIF3* expression in subcutaneous adipose tissue were obtained from GTEx Portal (https://www.gtexportal.org/home/) on 10/5/2022.

### Cell culture

A human white pre-adipocyte cell line (hWAs) was generated and kindly shared by Professor Yu-Hua Tseng (Joslin Diabetes Center, Harvard Medical School, USA) (41). For expansion, cells were cultured in 25 mM DMEM growth medium with GlutaMAX™ (10566016, ThermoFisher Scientific), 10% FBS (HyClone, GE Healthcare, Uppsala, Sweden) and 1% (100 U/ml) penicillin/streptomycin (15140122, Thermo Fisher Scientific). The cells were passaged at 90% confluence.

### DNA isolation and luciferase reporter assays

Genomic DNA was isolated from hWAs cells using DNeasy Blood and Tissue kit (69506, Qiagen) according to the manufacturer’s manual. To fine map the transcriptional regulatory regions in the *MTIF3* locus, we first identified the common genetic variants which were in tight linkage disequilibrium (r^2^ >= 0.8) with the lead variant rs1885988 in HaploReg v4.1(14). The thus identified 31 SNPs were tiled down into 12 DNA segments of the *MTIF3* gene, as shown in Supplemental Table 1. These segments were then PCR-amplified from hWAs DNA, all ranging from 700-1600 bp in size (depending on PCR primer design constraints), and with all SNP loci located several hundred bp from the ends of each fragment. The PCR primers were also designed to include flanking KpnI and EcoRV sites to allow cloning into the pGL4.23 minimal promoter luc2 luciferase reporter vector (Promega). For the reporter assays, hWAs were seeded into 96-well plates, and on the following day transfected with 95 ng of the pGL4.23 vectors and 5 ng pGL4.75 CMV-Renilla reporter vectors (for normalization), using Lipofectamine 3000 (ThermoFisher Scientific), in technical duplicates. Two days after transfection, the luc2 and Renilla signals were detected using Dual-Glo Stop&Glo reagents (E2920, Promega). The averages of technical duplicates were used to calculate luc2:Renilla ratios, which were then Z score-normalized to allow statistical evaluation across four independent experiments.

### gRNAs and ssDNA design for CRISPR/Cas9 mediated editing of rs67785913 in hWAs cells

To edit the rs67785913 CT allele to the minor CTCT allele in hWAs cell genome, CRISPR/Cas9 D10A nickase (Alt-R® S.p. Cas9 D10A Nickase V3, IDT) and two sgRNAs, and an ssDNA donor template were used. The sgRNA spacer sequences were: 5’-TTCAATAAGAAATTCCTCAA-3’ and 5’-GAAGAAAAAGGGGGGACACG-3’. The ssDNA sequence was 5’-TGTGGACTCGCAGTCTGCCCTTGAGGAATTTCTTATTGAAGAAGAAAAAGAGGGGGGACACGGGGCCCAGACCCCCAGCACCCGGCTTTCGAGCAGGCTC-3’. All oligonucleotides, sgRNAs, and ssDNA were purchased from Integrated DNA Technologies. The transfection was performed using Nucleofector 2b device (program A-033) (Lonza, Sweden) in nucleofector reagent L (Lonza, Sweden) mixed with 5 × 10^5^ hWAs cells, 120 pmol Cas9 nickase, 104 pmol sgRNA, and 300 pmol ssDNA. To increase the homology directed repair (HDR) editing efficiency, cells were incubated at 32 °C for 2 days in growth medium containing 30 μM HDR enhancer (Alt-R™ HDR Enhancer V2, IDT). Subsequently, cells were transferred to 37 °C for 3 days. For single-cell cloning, the hWAs cells were seeded at low density (2 cells/well in a 96-well plate) and allowed to expand for 3 weeks. Then the genomic DNA was extracted using QuickExtract DNA Extraction Solution (Lucigen) from the apparent single cell clonal populations. To identify the allele-edited homozygous clones, PCR was used to amplify the DNA fragment surrounding rs67785913 using primer pairs as below: Forward 5’-3’ GATTTGCAGGTGAGCAGACA, Reverse 5’-3’ ACTTGGAAATGGCCAAGATG; the amplicon was then subjected to Sanger sequencing to confirm the DNA sequence of each clone.

### Generation of inducible CRISPR/Cas9-expressing hWAs cell line (hWAs-iCas9)

hWAs cells were first seeded at 80,000 cells/well in six-well plates and transfected with 200 ng Super PiggyBac transposase (PB210PA-1, System Biosciences) and 500 ng pPB-rtTA-hCas9-puro-PB plasmid (kind gift from Dr. William Pu) (42) using Lipofectamine 3000 (ThermoFisher Scientific). The plasmid carries a doxycycline-inducible promoter driving the expression of Cas9, and a puromycin resistance gene, all flanked by piggyBac transposon integration sequences. After two days, the transfected cells were selected and expanded for three weeks in growth medium with 1 μg/ml puromycin, to obtain cells with genomically integrated inducible Cas9 construct.

### Differentiation of hWAs-iCas9 pre-adipocytes into mature adipocytes

hWAs-iCas9 pre-adipocytes were seeded into 24- or 96-well plates at the density of 40,000 or 8,000 cells per well respectively. After three days the cells reached confluency and were then incubated for 12 days with the differentiation cocktail, with medium changes every three days, as described before (41). To increase the accumulation of lipid droplets, 30 μM FFA (Linoleic Acid-Oleic Acid-Albumin) (L9655, Sigma-Aldrich) was added to the differentiation medium (43).

### CRISPR/Cas9 guide RNA design and off-targeting check

To generate *MTIF3* knock-out adipocytes, guide RNA spacer sequence targeting *MTIF3* exon 5, expressed in all known *MTIF3* protein-encoding transcripts (as reported at https://www.ensembl.org), was selected. The spacer sequence was 5’-GCAATAGGGGACAACTGTGC-3’, and full-length single guide RNA (sgRNA) was purchased from IDT. Furthermore, the hWAs genomic sequence surrounding the gRNA binding site was amplified by PCR using the primers 5’-CCACTTGTCTTGGGGACAGT-3’ and 5’-CTGGGAATGGTGGTTGAATC-3’, then analyzed by Sanger sequencing to ensure sequence match between gRNA spacer and the intended target locus. The potential off-target sites were predicted using CRISPR-Cas9 guide RNA design checker (https://eu.idtdna.com), and the genomic regions surrounding the top five off-target sites were PCR-amplified from the genomic DNA extracted from *MTIF3*-knockout and scramble control cells. The amplicons were then analyzed for any heteroduplexes generated by off-targeting using T7EI assay (IDT, Alt-R Genome Editing Detection Kit).

### sgRNA transfection and *MTIF3* knockout in mature adipocytes

After 12 days of differentiation, Cas9 expression was induced in mature adipocytes by adding 2 μg/ml doxycycline to the growth medium. On the following day, 30 nM pre-designed sgRNA was delivered into the cells using Lipofectamine RNAiMAX (13778075, ThermoFisher Scientific) according to manufacturer’s protocol; in parallel, 30 nM negative control crRNA (1072544, IDT) was used to transfect the scrambled control cells. One day post-transfection, cells were washed with PBS and incubated in normal growth medium for at least 3 days before functional assays carried out.

### Glucose restriction challenge for hWAs-iCas9 adipocytes

Mature hWAs-iCas9 adipocytes were firstly transfected with sgRNA against *MTIF3* to induce MTIF3 knockout or scramble crRNA to be as the controls described above. 3 days later, the mature adipocytes were incubated in DMEM medium (11966025, ThermoFisher Scientific) supplemented with different concentration of glucose for glucose restriction test (GR: 5 mM, 3 mM, and 1 mM) comparing with the growth condition *in vitro* (NF: 25 mM). The NF cells and GR cells were incubated for 3 days before the cell samples were collected and assays conducted.

### RNA isolation and qPCR gene expression assays

Total RNA was extracted from cells using RNeasy plus Kit (74034, QIAGEN) together with Qiazol reagent (79306, QIAGEN). RNA purity was assessed using Nanodrop (Nanodrop, Wilmington, USA), and cDNA was synthesized using SuperScript™ IV VILO™ Master Mix (11756500, ThermoFisher Scientific). Then, RT-qPCR was performed on ViiA7 qRT-PCR system (PE Applied Biosystems, Foster City, CA, USA), using predesigned Taqman assays following manufacturer’s instructions. The Taqman assays (ThermoFisher Scientific, Uppsala, Sweden) were: *MTIF3* (Hs00794538_m1), *GTF3A* (Hs00157851_m1), *TFAM* (Hs01073348_g1), *MT-CO1* (Hs02596864_g1), *PRDM16* (Hs00223161_m1), *TOMM20* (Hs03276810_g1), *CPT1B* (Hs00189258_m1), *ACADM (Hs00936584_m1), ACAT1* (Hs00608002_m1), *ABHD5* (Hs01104373_m1), *PNP1A2* (Hs00386101_m1), *ACACB* (Hs01565914_m1), *MT-ND1* (Hs02596873_s1), *MT-ND2* (Hs02596874_g1), *MT-ND3* (Hs02596875_s1), *MT-ND4* (Hs02596876_g1), *MT-CO2* (Hs02596865_g1), *MT-CO3* (Hs02596866_g1), *HPRT-1* (Hs99999909_m1), *TBP* (Hs00427620_m1) and *RPL13A* (Hs03043885_g1). The relative gene expression was calculated using the delta Ct method, and the target gene expression was normalised to the mean Ct of three reference genes *HPRT-1, TBP* and *RPL13A*.

### Western blotting

Cells were washed twice with ice-cold PBS and lysed in 1% SDS buffer for 10 mins, then passed through a QIAshredder (79654, Qiagen) and centrifuged for 15 mins at 14,000 x g. The supernatant was subsequently collected and protein concentration quantified using the BCA assays (23225, ThermoFisher Scientific). To assess target protein expression, 10 μg lysates were loaded into 4–20% Mini-PROTEAN® TGX Stain-Free™ Protein Gels (Bio Rad Laboratories AB, Solna, Sweden) and separated, followed by transfer of polyvinylidene difluoride (PVDF) membranes (1704156, Bio Rad Laboratories AB). After blocking in 5% BSA solution for 1 hour, the membranes were incubated with primary antibodies against MTIF3 (14219-1-AP, Proteintech), OXPHOS complex (45-8199, ThermoFisher Scientific), FABP4, ACC, FAS (12589, Cell Signalling Technology) and corresponding HRP-conjugated secondary antibodies (anti-mouse IgG, Cell Signalling Technology; anti-rabbit IgG, Cell Signaling Technology). TBS with 0.1% (v/v) Tween-20 was used for membrane washing, and TBS with 2% BSA was used for antibody incubation. To visualize the blots, Clarity western ECL substrate was added to the membrane and a CCD camera used to acquire images and Image Lab software (Bio Rad Laboratories AB, Solna, Sweden) were used to develop the images. Image J software was used to quantify the protein bands. After detection of the protein targets, the membranes were stripped using Restore Western Blot Stripping Buffer (21059, ThermoFisher Scientific) and blotted using anti-β-Actin antibody (4967, Cell Signaling Technology) or anti-GAPDH antibody (ab37168, Abcam). Source data files are archived in Figure 3-source data.zip file.

### Mitochondrial function in *MTIF3* knockout adipocytes

To directly assess the effects of *MTIF3* on mitochondrial respiration in adipocytes we used the Seahorse XF (Seahorse Bioscience, North Billerica, MA) to measure cellular respiration oxygen consumption rate (OCR) under different conditions. hWAs-iCas9 cells were seeded at 8,000 cells per well in a Seahorse 24-well plate, then differentiated and induced for *MTIF3* knockout or with scrambled control, as described above. Then, cells were adapted in 1 g/L growth medium (31885049, ThermoFisher Scientific) for three days. Mitochondrial function was then assessed using the Seahorse XF-24 instrument according to a protocol optimized for the adipocyte cell line. Briefly, to measure OCR independent of oxidative phosphorylation, 2 μM oligomycin (O4876, Sigma-Aldrich) was added to the cells. Subsequently, 2 μM FCCP (carbonyl cyanide-p-trifluoromethoxyphenylhydrazone) (C2920, Sigma-Aldrich) and 5 μM respiratory chain inhibitors: rotenone (R8875, Sigma-Aldrich) and antimycin A (A8674, Sigma-Aldrich) were added to measure maximal respiration and basal rates of non-mitochondrial respiration. Cells were then frozen at −80 °C for at least 4 hours, then the plate was dried, and DNA was extracted with CyQUANT™ Cell Lysis Buffer (C7027, ThermoFisher Scientific). Total DNA was then quantified by Quant-iT™ PicoGreen™ dsDNA Assay Kit (P7589, ThermoFisher Scientific) against a lambda DNA-generated standard curve.

### Endogenous long chain fatty acid oxidation in adipocytes

The Seahorse mitochondrial analyzer was used to test the effects of MTIF3 loss on endogenous long chain fatty acid oxidation in adipocytes. Prior to the assay, adipocytes were incubated overnight with substrate-limited medium: DMEM (A14430, ThermoFisher Scientific); 0.5 mM glucose (103577-100, Angilent); 1.0 mM glutamine (103579-100, Angilent); 0.5 mM carnitine (C0283, Sigma-Aldrich); 1% FBS (SV30160.03, HyClone). On day of the assay, the substrate-limited medium was replaced with FAO assay medium: 1xKHB buffer was supplemented with 2.5 mM glucose, 0.5 mM carnitine and 5 mM HEPES, and the pH was adjusted to pH 7.4 with NaOH. The cells were then treated for 15 min with either 40 μM etomoxir (E1905, Sigma-Aldrich) or only with the solvent (DMSO). Etomoxir inhibits carnitine palmitoyltransferase (CPT)-1 and diglyceride acyltransferase (DGAT) activity in mitochondria, and thus inhibits mitochondrial fatty acid oxidation (44, 45). The OCR was then measured as described above.

### Mass spectrometry-based metabolite profiling

The mature hWAs-iCas9 adipocytes were firstly induced for *MTIF3* knockout, followed by glucose restriction challenge as described above. The cells were quenched on dry ice and metabolites were extracted using a previously optimized protocol (46).

For analysis of low molecular weight metabolites, extracts were reconstituted in 100 μl of MeOH/water (8/2, v/v) and 60 μl was transferred to new Eppendorf tubes and evaporated to dryness using a miVac concentrator (SP Scientific, NY) for 3 h at 30 °C. Dried samples were methoximated using 20 μl of methoxyamine hydrochloride in pyridine (Thermo Scientific, MA) by shaking at 3000 rpm for 30 min at room temperature (VWR, PA). Afterward, 20 μl of N-methyl-N-(trimethylsilyl) trifluoroacetamide (MSTFA) +1% trimethylsilyl chloride (Thermo Scientific, MA) was added to each sample and shaken at 3000 rpm at room temperature for 1 h. Samples were transferred to glass vials and immediately analyzed using an Agilent 6890 gas chromatograph connected to an Agilent 5975CL VL MSD mass spectrometer controlled by MassHunter Workstation software 10.0 (Agilent, Atlanta, GA). One μl sample was injected at 270 °C on an HP-5MS column (30 m length, 250 μm ID, 0.25 μm phase thickness). with a helium gas flow rate of 1 ml/min and a temperature gradient starting at 70 °C for 2 min, increasing 15 °C/min to 320 °C and held for 2 min. Data were acquired using electron ionization (EI) at 70 eV in either full scan (50-550 *m/z*) or single ion monitoring (SIM) mode. The MS-DIAL version 4.7 was used for raw peak extraction, peak alignment, deconvolution, peak annotation, and integration of peaks.

Amino acids and free fatty acids were chemically derivatized and analyzed using a previously described method (47). Briefly, 40 μl of the samples were mixed with 20 μl of 3-nitrophenylhydrazine (3-NPH) (Sigma-Aldrich, MO), followed by addition of 20 μl of 1-ethyl-3-(3-dimethylaminopropyl)carbodiimide hydrochloride (EDC) (Merck, Millipore, MO) and shaking at 3000 rpm at room temperature for 1 h. Samples were analyzed using an Agilent 1260 ultra-performance liquid chromatograph coupled with an Agilent 6495 tandem mass spectrometer and controlled by MassHunter version 8.0 (Agilent Technologies, CA). Three μl sample was injected on an Agilent Eclipse RRHD C18 column (2.1 × 150 mm, 1.8 μm) (Agilent Technologies, CA) with a flow rate of 0.6 ml/min and a column oven temperature of 50 °C. The mobile phase A and B were 0.1% formic acid (Fisher Chemical, Prague, Czech Republic) in Milli-Q water (Merck, Millipore, MO) and acetonitrile (VWR, Paris, France), respectively. Gradient elution was performed as follows: held at 5% B from 0–1 min, changed linearly to 90% B in 10 min, changed from 90% B to 100% B in 13 min, held at 100% B for 2 min, returned to 5% B (initial condition) in 0.1 min, and held at 5% B for 2 min. Analyses were conducted in negative electrospray ionization mode (ESI) mode with the nebulizer gas pressure set at 20 psi, ion capillary voltage at 2500 V, gas temperature at 150 °C, and sheath gas temperature at 250 °C. Data were recorded in multiple reaction monitoring (MRM) mode, with two transitions for each analyte.

### Lipolysis quantification in differentiated hWAs cells

Differentiated scrambled control or *MTIF3* knockout cells were washed twice with PBS and then incubated with DMEM containing 2% free fatty acid-free BSA for 2 hours. For the insulin or isoproterenol-stimulated lipolysis, 100 nM insulin (I2643, Sigma-Aldrich) or 10 μM isoproterenol (1351005, Sigma-Aldrich) was added in the medium separately. After the incubation, the medium was collected, and the glycerol content was measured using Glycerol-Glo™ Assay (J3150, Promega).

### Total triglycerides measurement

Triglyceride-Glo™ Assay kit (J3161, Promega) was used to quantify total triglyceride content in scrambled control or *MTIF3* knockout cells before and after glucose restriction treatment and after refeeding. Briefly, cells were collected in 50 μl kit lysis buffer at room temperature for 1 hour. Then 2 μl lysate was mixed with 8 μl glycerol lysis solution with lipase, and incubated at 37 °C for 30 min. Subsequently, 10 μl glycerol solution was mixed with 10 μl glycerol detection solution supplemented with reductase substrate and kinetic enhancer, and transferred into a 384-well plate. After 1 h incubation at room temperature, the luminescence was detected using CLARIOstar plate reader (BMG Labtech, Germany), and the triglyceride concentration was calculated using a standard curve generated from glycerol standards and normalized to total protein measured using BCA assays (23227, ThermoFisher Scientific).

### Data and Resource Availability

All data and resource generated in this study are available upon reasonable request.

### Statistics

For each assay, the number of biological and technical replicates, standard deviation and statistical significance are reported in the figure legends. Hypothesis tests were performed using two-tailed Student’s t-test, one-way ANOVA or paired t-test. A nominal p-value of <0.05 was considered statistically significant. All analyses were undertaken using Prism GraphPad 9.0 software (La Jolla California, USA), SIMCA 17.0 (Sartorius Stedim Data Analytics, Malmö, Sweden), Rstudio 1.4, and Microsoft Excel 365. For the metabolome data, ANOVA (aov) was performed in R with genotype and feeding condition as independent variables with Tukey’s test post hoc (TukeyHSD). Significance was defined as q<0.05 using multiple testing adjustment according to the false discovery rate (p.adjust).

## Acknowledgements

We thank Jennifer Doudna (UC Berkeley) for initial support with concepts relating to using CRISPR in *in vitro* studies of gene-environment interactions. We also thank Yu-Hua Tseng in Joslin Diabetes Centre for providing the hWAs cells.

This work was supported by China Scholarship Council (201708420158 to Mi Huang); the European Commission (ERC-2015-CoG -681742 NASCENT to PWF), and Swedish Research Council (Distinguish Young Research Reward in Medicine) (to PWF), LUDC-IRC and The Swedish Research Council (to HM), and by The Albert Påhlsson Foundation (to SK).

---

**Supplemental Table 1.** 31 SNPs in tight linkage disequilibrium (r2 >= 0.8) with the lead variant rs1885988 tiled down into 12 DNA segments of the *MTIF3* gene for luciferase reporter assay

| Rs ID | Position at hg38 chr13 | Luciferase reporter construct |
| --- | --- | --- |
| rs7988412 | 27426145 | 1 |
| rs76790205 | 27428748 | 2 |
| rs10220056 | 27429644 |  |
| rs9581848 | 27429860 |  |
| rs9581849 | 27431645 | 3 |
| rs74183666 | 27433628 | 4 |
| rs7669 | 27435714 | 5 |
| rs1885989 | 27435980 | 6 |
| rs1885988 | 27436125 |  |
| rs57724994 | 27436252 |  |
| rs76249173 | 27436308 |  |
| rs75439483 | 27436375 |  |
| rs77055070 | 27436379 |  |
| rs143190464 | 27436414 |  |
| rs74563672 | 27436523 |  |
| rs147240135 | 27436782 |  |
| rs12867531 | 27436791 |  |
| rs78334317 | 27436817 |  |
| rs76488435 | 27436853 |  |
| rs79023532 | 27436942 |  |
| rs74811644 | 27437021 |  |
| rs45622135 | 27437457 | 7 |
| rs9581850 | 27438128 |  |
| rs12018313 | 27438129 |  |
| rs139686876 | 27438919 | 8 |
| rs9581852 | 27439611 |  |
| rs9579083 | 27443133 | 9 |
| rs9581854 | 27443645 | 10 |
| rs9581855 | 27443877 |  |
| rs67785913 | 27451171 | 11 |
| rs9512699 | 27455759 | 12 |

**Supplemental Table 2.**
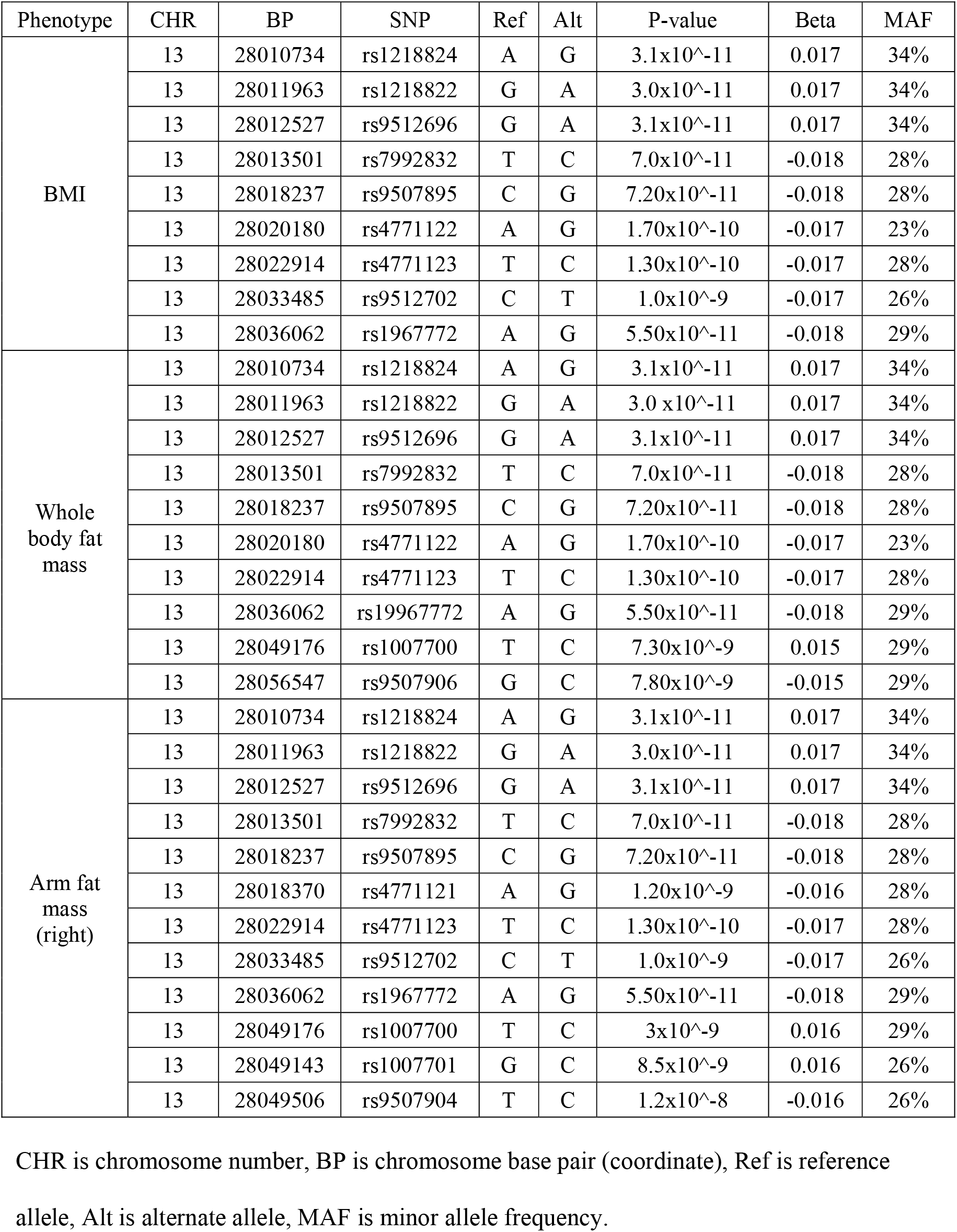
SNPs in *MTIF3* locus nominally associated with BMI, whole body fat mass and arm fat mass (right)

